# Ecologically inspired assembly of a *Bacillales* synthetic community for lignin degradation

**DOI:** 10.64898/2026.08.24.746789

**Authors:** Qing Zhou, Xingchen Xin, Chao Du, Youri Meelkop, Sebastiaan N. J. Laan, Joost Willemse, Imke Bonné, Maaike van Geel, Gilles P. van Wezel, Ákos T. Kovács, Johannes H. de Winde, Xinming Xu

**Author notes:** Correspondence (Á.T. Kovács), (Winde, J.H. de), (X. Xu).

## Abstract

Lignin remains a major bottleneck in biomass valorization due to its heterogeneous and recalcitrant structure. Traditionally, lignin degradation has been developed using a single-microorganism or single-enzyme approach, with efforts focused on identifying ligninolytic activities of individual bacteria or fungi. Here, we establish a framework for designing lignin-degrading synthetic microbial communities (SynComs) by integrating top-down ecological filtering with bottom-up trait-based selection. This approach enables the selection of bacteria with ligninolytic capacity while retaining community members that form strong biofilms and contribute indirectly to overall community function. Proteomic profiling uncovered lignin-driven functional reprogramming of the SynCom, highlighting strain-specific enzymatic specializations that collectively enable lignin depolymerization. Our work highlights the potential of SynComs for lignin degradation and shows how rationally designed, functionally partitioned communities can enhance lignin conversion.

## Introduction

Lignocellulose, composed mainly of cellulose, hemicellulose and lignin, is the primary structural component of plant cell walls (Rinaldi et al., 2016). Lignocellulose is the most abundant renewable carbon resource on Earth (Bomble et al., 2017). Among its three major components, lignin is a heterogeneous aromatic polymer composed of p-hydroxyphenyl (H), guaiacyl (G) and syringyl (S) units that are linked by various covalent bonds, primarily ether and carbon–carbon linkages (Ralph et al., 2004). This complex organization of lignin polymers is a major factor resulting in their high recalcitrance and resistance to biodegradation (Lankiewicz et al., 2023). Although lignocellulose is an attractive renewable carbon source and lignin is a rich reservoir of aromatic compounds, the lack of efficient lignin-degrading technologies limits its efficient utilization and largely relegates lignin to low-value combustion (Ragauskas et al., 2014). Therefore, developing environmentally friendly and sustainable strategies for lignin depolymerization is crucial for unlocking its full valorization potential.

Biological approaches to lignin degradation have gained increasing momentum in recent years. Microorganisms such as fungi and bacteria have been demonstrated to degrade lignin efficiently and sustainably (Atiwesh et al., 2022). Fungi have long been recognized as efficient lignin degraders, and several white-rot and filamentous species (e.g. *Aspergillus*, *Pleurotus*, *Lentinus*, *Trametes*) can remove more than 50–70% of lignin from lignocellulosic feedstock within a week or two under optimized conditions (Janusz et al., 2017; Virmani et al., 2024; Wu et al., 2005). However, their strict requirements for aeration, moisture, and substrate conditions often limit large-scale industrial applications. In contrast, performance of bacteria is generally more robust, tolerating wider ranges of temperature, pH and oxygen, which makes them attractive chassis for industrial lignin valorization (R. Xu et al., 2018). Diverse bacterial genera isolated from soil, decaying wood, pulp, paper wastewater, and animal guts have been reported to degrade lignin, mainly via secretion of extracellular peroxidases and laccases that cleave key linkages in the polymer (Bugg, 2024).

Among lignin-degrading bacteria, *Bacilli* stand out as widespread soil bacteria with remarkable robustness and biotechnological potential. Several isolates of the *Bacillales* order have been reported to degrade lignin across a variety of substrates, including alkaline and kraft lignin, straw lignin, and pulp-and-paper effluents. For instance, *Bacillus altitudinis* SL7 was reported to remove up to 44% of alkali lignin, with extracellular laccase activity hypothesized to play a central role in catalyzing oxidative depolymerization (Khan et al., 2021). Likewise, a *Bacillus velezensis* strain demonstrated 40% lignin degradation under optimized conditions and was shown to produce multiple ligninolytic enzymes, including lignin peroxidase (LiP), manganese-dependent peroxidase (MnP), and laccase (Verma et al., 2020). Beyond their hydrolase repertoire, many *Bacillales* strains synthesize abundant extracellular polymeric substances (EPS) that encapsulate cells and promote adhesion and aggregation on surfaces (Yadav et al., 2023). Biofilm formation is widely recognized to enhance polymer biodegradation by facilitating close cell–substrate contact and strengthening microbial and enzymatic interactions, as demonstrated in plastic-degrading systems (Chattopadhyay, 2022). Within biofilms, the EPS matrix functions as a ‘digestive system’ by retaining extracellular enzymes in close proximity to the substrate (McFall et al., 2024). Consequently, enhanced biofilm formation in plastic-degrading bacteria has been shown to accelerate overall biodegradation rates (S. Li et al., 2025). While this mechanism has been documented in plastics biodegradation, the potential role of biofilm formation in lignin degradation has been largely overlooked. Collectively, these observations suggest that, in addition to their enzymatic potential, the biofilm-forming capacity of *Bacillales* may represent an important and underexplored determinant contributing to the efficiency and stability of lignin biodegradation.

In nature, wood decomposition is inherently a community-driven process; it is never carried out by a single bacterium or fungus but by the collective, synergistic actions of diverse microbial guilds. Building on this natural division of labor, some studies have combined functionally distinct microorganisms, using fungi act as ‘pioneers’ that deploy oxidative enzymes and cellulases to breach the recalcitrant lignocellulosic matrix, and bacteria as ‘workhorses’ that secrete thermostable hydrolases to further deconstruct the exposed polymers (Ding et al., 2025). In addition, some yeast strains, although not exhibiting substantial degradation activities, were incorporated into lignocellulose-degrading consortia because their production of metabolites derived from released carbohydrates, supplemented the activities of other community members (Vu et al., 2023). Thus, artificially assembled microbial consortia with functional partitions provide a novel approach for lignin degradation. In recent years, synthetic microbial communities (SynComs) have been widely applied in plant growth promotion (Xu et al., 2025), environmental remediation and gut microbiota engineering, yet, they are rarely explored in the context of lignin degradation. Typically, SynCom assembly is classified into bottom-up or top-down approaches. The former begins with selecting individual strains as functional building blocks and assembling them into a community to achieve emergent properties unattainable by single strain. The latter starts from a complex community and subsequent identification of keystone members while filtering out functionally redundant taxa, with the goal of recapitulating the original functionality using a simplified subset of microbes. Both approaches have their own advantages and limitations. Bottom-up approaches are typically labor-intensive, requiring extensive screening and selection of strains with desired functions, but may omit taxa that are essential for community performance while not directly contributing to the focal function. In contrast, top-down approaches often rely on metataxonomics and tend to infer causality between taxon prevalence and functional activity, potentially ignoring low-abundance taxa that may exhibit high functional performance.

Here, we integrate bottom-up and top-down strategies to construct a lignin-degrading SynCom, thereby overcoming the limitations of relying on either approach alone. We hypothesize that an effective bacterial consortium combines robust biofilm formers and strains possessing ligninolytic capacity. Our integrative design enables the retention of taxa that may not directly contribute to lignin degradation but are ecologically relevant, while effectively selecting and preserving key functional degraders within the community.

## Methods

### Bacterial strains and growth media

121 strains used in this work are from the order *Bacillales* (Supplementary Table 1) (Song et al., 2024). Lysogeny broth (LB, Lennox, Carl Roth, 10 g/L tryptone, 5 g/L yeast extract, and 5 g/L NaCl) or LB agar media were used for cultivating *Bacillales* strains. Beechwood lignin (average molecular weight, 2900 g/mol) was extracted at pilot-scale (Fraunhofer CBP) using the Fabiola aqueous acetone organosolv process from Netherlands Organisation for Applied Scientific Research (TNO) and dissolved in dimethyl sulfoxide (DMSO) at 50 g/L as a stock solution. M9 minimal medium was used to provide a nutrient-limited environment and promote the utilization of lignin as the main carbon source. The M9 base contained 6.78 g/L Na□HPO□, 3.0 g/L KH□PO□, 0.5 g/L NaCl, and 1.0 g/L NH□Cl. The lignin stock was added to the medium to obtain a final lignin concentration of 1 g/L. After autoclaving, the medium was supplemented with 2 mL/L of 1 M MgSO (final concentration 0.05 g/L), 100 μL/L of 1 M CaCl□ (final concentration 0.002 g/L), and 20 mL/L of a 25% (w/v) glucose solution. For M9 agar, 1.4% (w/v) agar was added. For liquid cultivation, 0.2% of glucose was included to support initial bacterial growth.

### Screening single strains for lignin degradation

A total of 121 strains from the *Bacillales* collection were arrayed in flat-bottom 96-well plates (Sarstedt) as glycerol stocks by mixing overnight LB cultures with glycerol to a final concentration of 40% (v/v) in a total volume of 100 μL per well. Glycerol stocks were stamped onto M9–lignin agar plates using a 96-pin steel stamp, and the plates were incubated at 30 °C.

### Identification and comparative analysis of mhq-associated gene clusters

Homologous genes of the *mhq* operon (*mhqD*/*O*/*P*) were searched against *Paenibacillus* genomes using cblaster (v1.4.0) in local mode with DIAMOND databases (Buchfink et al., 2015; Gilchrist et al., 2021). Genomic contexts were extracted from the resulting session files and visualized using clinker for synteny comparison. Cluster boundaries were restricted to ±20 kb around query hits, and representative clusters containing ≥2 *mhq* homologs were used for final visualization.

### Isolation and Identification of lignin-associated strains

A total of 121 *Bacillales* strains from the collection were first grown overnight in LB liquid individually. The cultures were adjusted to an OD□□□ of 0.80–0.85, and 10 μL of each adjusted culture was inoculated into M9-lignin medium. All cultures were incubated in flasks at 30 °C using 200 rpm shaking. The cultivation of 121 *Bacillales* strains in M9-lignin medium resulted in a ring-shaped biofilm along the inner wall of the flasks. The ring-shaped biofilm was collected after 7 days and plated in serial dilutions onto LB agar medium to isolate bacteria associated with lignin degradation. Phenotypically distinct colonies were clean streaked for purity. DNA of isolates was extracted by lysis of 30 μL of bacterial o/n cultures in 50 μL of buffer I containing 25 mmol/L NaOH, 0.2 mmol/L EDTA, pH 12 at 95 °C for 30 min, before the pH value was lowered by addition of 50 μL of buffer II containing 40 mmol/L Tris-HCl at pH 7.5. To specifically identify and differentiate the *Bacillales* species, the *tuf* gene was amplified using the primer pair tuf1-F (5′-CACGTTGACCAYGGTAAAACH-3′) and tuf2-R (5′-GTDAYRTCHGWWGTACGGA-3′), which target a 982 bp region. Each 25 μL PCR reaction contained 12.5 μL TEMPase Hot Start 2× Master Mix Blue, 0.8 μL of each primer (10 μmol/L), 10.6 μL nuclease-free water, and 0.3 μL DNA template. The thermal cycling program consisted of an initial denaturation at 95 °C for 15 min; 30 cycles of 95 °C for 30 s, 47 °C for 30 s, and 72 °C for 1 min; followed by a final extension at 72 °C for 5 min. PCR products were purified with the NucleoSpin Gel and PCR Cleanup Kit (Macherey–Nagel) and subsequently sequenced by Eurofins Genomics. The resulting sequences were blasted at the NCBI Nucleotide BLAST database (Dataset S1), specifically the WGS contigs under BioProject ID 960711, enabling precise identification of each isolate in our *Bacillales* library.

### DNA extraction and Oxford Nanopore sequencing

To assess community-level associations with lignin-induced biofilm formation, total genomic DNA was extracted from the complete ring-shaped biofilm using the DNeasy PowerSoil Kit (Qiagen, Hilden, Germany) according to the manufacturer’s protocol. The *tuf* primer pair described above was used for Oxford Nanopore sequencing. DNA concentration and purity were assessed using a NanoDrop spectrophotometer, and DNA quantity was further determined with the Quant-iT dsDNA High-Sensitivity Assay Kit (Thermo Fisher Scientific).

For Oxford Nanopore sequencing, libraries were prepared using the SQK-RBK114.96 Rapid Barcoding Kit (Oxford Nanopore Technologies, Oxford, UK), starting with 200 ng of genomic DNA per sample. Briefly, each sample was simultaneously fragmented and barcoded using a unique rapid barcode, with incubations at 30 °C for 2 min and 80 °C for 2 min. Barcoded samples were pooled in equimolar ratios and purified with AMPure XP beads (AXP). Subsequently, 1 μl Rapid Adapter (RA) was ligated to 11 μl of the pooled library, and the final library concentration was measured using NanoDrop. Prepared libraries were loaded onto a Flongle flow cell (FLO-FLG114) and sequenced on a MinION Mk1B device following the standard ONT protocol. Raw reads were base called and demultiplexed using MinKNOW v25.05.14. Raw reads sequencing on Nanopore have been deposited in NCBI Bioproject under PRJNA1505615.

We established an amplicon analysis workflow adapted from ONT-AmpSeq to process the Nanopore sequencing data (https://github.com/Xinming9606/ONT_Amp_analysis) (Schacksen et al., 2024). Briefly, Nanoplot was used to generate read statistics and quality plots for each barcode, providing an overview of read quality and length distribution (De Coster and Rademakers, 2023). Amplicons were filtered and trimmed with Chopper (parameter: -l 400, -u 1200, -q 20) (De Coster and Rademakers, 2023). Draft consensus sequences were generated with VSEARCH (Rognes et al., 2016), and mapped using Minimap2 and polished with Racon (https://github.com/lbcb-sci/racon) to obtain high-quality consensus OTUs (Li, 2018). Final OTU tables were constructed by VSEARCH clustering at 99% sequence identity. Taxonomy was assigned using BLAST+ against the *Bacillales tuf* gene reference database (https://github.com/Xinming9606/KovacsLab-BLASTdb).

### Biofilm formation assay

The *Bacillales* strains were screened for their biofilm formation ability using a crystal violet staining assay (Yadav et al., 2023). 1 μL of overnight culture was inoculated into 200 μL tryptic soy broth (TSB, Merck Millipore, 17 g/L casein peptone, 2.5 g/L dipotassium hydrogen phosphate, 2.5 g/L D(+)-glucose, 5 g/L sodium chloride, and 3 g/L soy peptone) in 96-well plates and incubated at 37 °C for 24 h without agitation. Wells containing 200 μL TSB without inoculum served as negative controls. The culture was then removed, and wells were washed thrice with phosphate buffer saline (PBS, pH = 7.2). The attached biofilm was fixed for 10 min with 200 μL methanol, air-dried for 30 min, and then stained for 10–15 min with 200 μL crystal violet (1%, w/v). The unattached stain was removed by washing with PBS and air-dried for 30 min in an inverted position. The attached dye was removed by the addition of 200 μL of 33% v/v glacial acetic acid. Absorbance was measured at 620 nm using TECAN Spark 10M multimode microplate reader. All assays were performed in triplicate wells for each strain.

### Dye decolorization assay

Lignin-mimicking dyes, methylene blue (MB) and Azure B (AB) were used for ligninolytic activity assays. The decolorization ability of *Bacillales* strains was determined using a quantitative dye decolorization assay, as described by Bharti et al. with modifications (Bharti et al., 2019). Overnight cultures were inoculated into fresh LB until the OD_600_ reached 1.0. Aliquots of 200 µL were transferred into 96-well plates supplied with 50 mg/L MB or AB and incubated at 30 °C for 72 h. Following incubation, the culture was centrifuged, and the absorbance of the supernatant was measured at 665 nm (λmax of MB) or 650 nm (λmax of AB) using TECAN Spark 10M multimode microplate reader. Cell-free incubations were assayed as control. The decolorization ratio was measured according to the following equation:

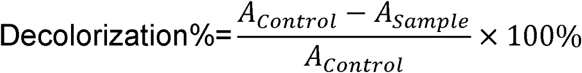

All the experiments were conducted in triplicate.

### Scanning Electron Microscopy (SEM)

SEM imaging was conducted to analyze the biofilm formation on the surface of lignin. The lignin degrading SynCom (LDSynCom) was cultivated in M9 minimal medium containing 1 g/L lignin and 0.2% (w/v) glucose for 10 days. After incubation, the ring-shaped biofilm was carefully collected and used as samples. Lignin incubated in medium without the LDSynCom served as an abiotic control. The samples were fixed with 1.5% glutaraldehyde (30 min). Subsequently, samples were dehydrated using a series of increasing ethanol percentages (70%, 80%, 90%, 96%, and 100%, each step 30 min) and critical point dried (Baltec CPD-030). Subsequently, the samples were coated with 10 nm Platinum palladium using a sputter coater and directly imaged using a JEOL JSM6700F. Images were acquired with 5kV at a working distance of 8mm.

### SynCom Lignin degradation assay

The LDSynCom was tested for lignin degradation. Individual strains were grown overnight in LB, adjusted to an OD of 0.8, and 100 μL of each culture was inoculated into M9-lignin medium shake flasks. Cultures were incubated at 30 °C with shaking, and at the indicated time points OD was measured. The absorbance at 280 nm was determined to characterize the lignin degradation (Z. Xu et al., 2018a). An uninoculated medium was used as the control.

Folin-Ciocalteu assay (FCA) was applied for detecting released small molecule phenolic products (Rashid and Bugg, 2021). All the experiments were conducted in triplicates. 20 μL of samples were mixed with 100 μL distilled water. And 10 μL Folin-Ciocalteu reagent was added, 100 μL of Na_2_CO_3_ solution (4%) was added after 4 min. Then incubated for 30 min in the dark at room temperature. Absorbance was measured at 750nm using TECAN Spark 10M multimode microplate reader.

### Protein extraction, Liquid Chromatography–Tandem Mass Spectrometry and statistics analysis

The LDSyncom was inoculated into M9 medium and M9 supplemented with lignin, with four biological replicates per condition. Cultures were incubated at 30 °C with shaking for 7 days. Samples were prepared as described with some modification (Gubbens et al., 2012; Zhang et al., 2020). Cultures were centrifuged to remove bacterial cells, and the cell-free supernatants were collected. Proteins in the supernatants were concentrated using 10 kDa Amicon Ultra-2 centrifugal filter devices (10 kDa MWCO, Millipore). Concentrated proteins were precipitated using the chloroform-methanol method and redissolved in 1% sodium deoxycholate at 95 °C. The protein concentration was measured at this step using BCA method. Protein samples were then reduced by adding 5 mM DTT and incubated at 60°C for 30 min, followed by thiol group protection with 21.6 mM iodoacetamide incubation at room temperature in dark for 30 min. Then 0.1 μg trypsin (recombinant, proteomics grade, Roche) per 10 μg protein was added, and samples were digested at 37°C for 8 h and then at room temperature. After digestion, trifluoroacetic acid was added to 1% and samples were incubated at 37°C for 30 min followed by centrifugation to precipitate deoxycholate. Peptide solution containing 6 μg peptide was then cleaned and desalted using STAGE-Tips (Rappsilber et al., 2007). Briefly, 6 μg of peptide was loaded on a conditioned StageTip with C18 disk (AttractSPE Tips C18.T1.10.960, Affinisep), washed once with 0.5% formic acid solution, and eluted with elution solution (80% acetonitrile, 0.5% formic acid). Acetonitrile was then evaporated in a SpeedVac. Final peptide concentration was adjusted to 40 ng·μL-1 using sample solution (3% acetonitrile, 0.5% formic acid) for analysis.

Peptides were analyzed on a Bruker timsTOF HT platform using diaPASEF. **LC** (**Liquid Chromatography**) separation was performed with a 55-min gradient at 0.3 μL min^-1 and 50 °C column temperature. Acquisition ranges were m/z 100–1700 and ion mobility 1/K0 0.70–1.35, with one MS1 frame followed by 10 DIA frames per cycle.

Raw data were processed in DIA-NN v2.2.0 in library-free mode against a protein database derived from the complete genomes of the SynCom members (Demichev et al., 2020). Core DIA-NN settings included –qvalue 0.01, –fasta-search, –predictor, – gen-spec-lib, –matrices, –reanalyse, one missed cleavage, peptide length 7–30 aa, precursor charge 2–4, precursor m/z 300–1800, and fragment m/z 200–1800. The in silico digest used –cut K*,R* without enforcing the proline rule.

MaxLFQ-aggregated protein intensities were normalized using Variance Stabilizing Normalization (vsn), followed by Perseus-style imputation of missing values. Differential protein abundance analysis was performed using limma with eBayes(trend = TRUE, robust = TRUE) and Benjamini–Hochberg correction (Benjamini and Hochberg, 1995). Proteins were considered significant at adjusted p < 0.05 and |log2FC| > 1. Functional enrichment was performed using Fisher’s exact test with FDR correction (Benjamini and Hochberg, 1995). Lignin-related candidates were identified by curated keyword matching against RefSeq descriptions plus EC number matching, and were further evaluated by hmmscan/Pfam domain evidence (Eddy, 2011; Mistry et al., 2021). Shared peptides were resolved using an EM-based strain deconvolution framework (Dempster et al., 1977). All analysis code is available at: (https://github.com/Xinming9606/lignin_SynCom_proteomics)

## Results

### Bottom-up selection of lignin degrading *Bacillales* strains

To rationally construct a lignin-degrading SynCom, we combined complementary bottom-up and top-down selection strategies to identify *Bacillales* strains with ligninolytic potential (Figure 1). In this bottom-up selection step, the whole 121-strain *Bacillales* library was assessed for lignin utilization by screening for clear halo formation around single strain colonies on M9 agar plates supplemented with different carbon sources, namely glucose, kraft lignin, and beechwood lignin (Figure 2A). Most strains produced colonies on glucose-containing medium after 7 days, whereas growth of most colonies was limited on lignin-containing medium. Of the 121 strains examined, *Paenibacillus* sp. 11B was the sole isolate that displayed clear halo formation on M9 agar medium supplemented with beechwood lignin (Figure 2B), pointing to lignin degradation or modification activity of the strain.

**Figure 1.**
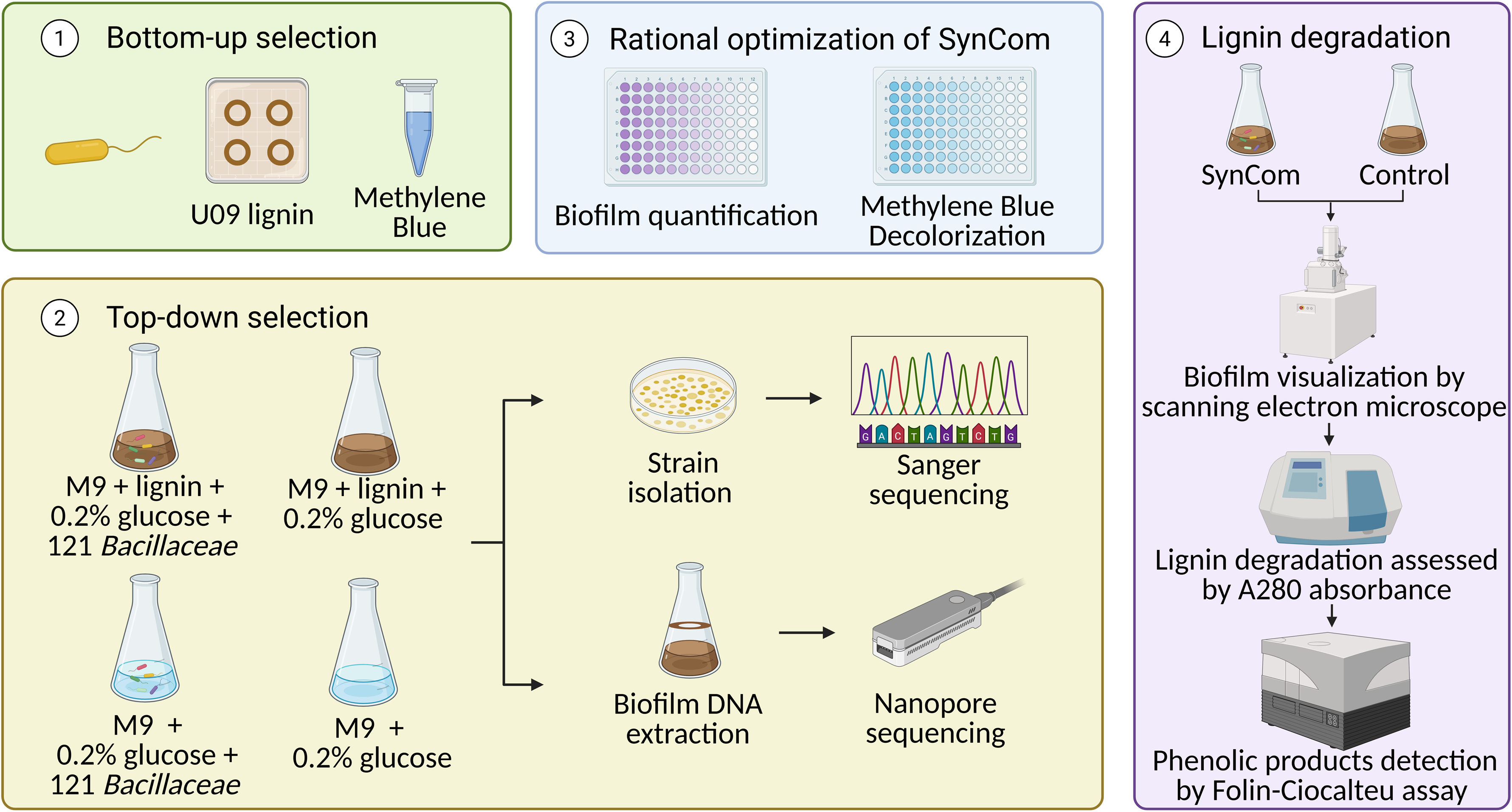
Schematic overview of the integrated workflow for rational SynCom design and optimization.

**Figure 2.**
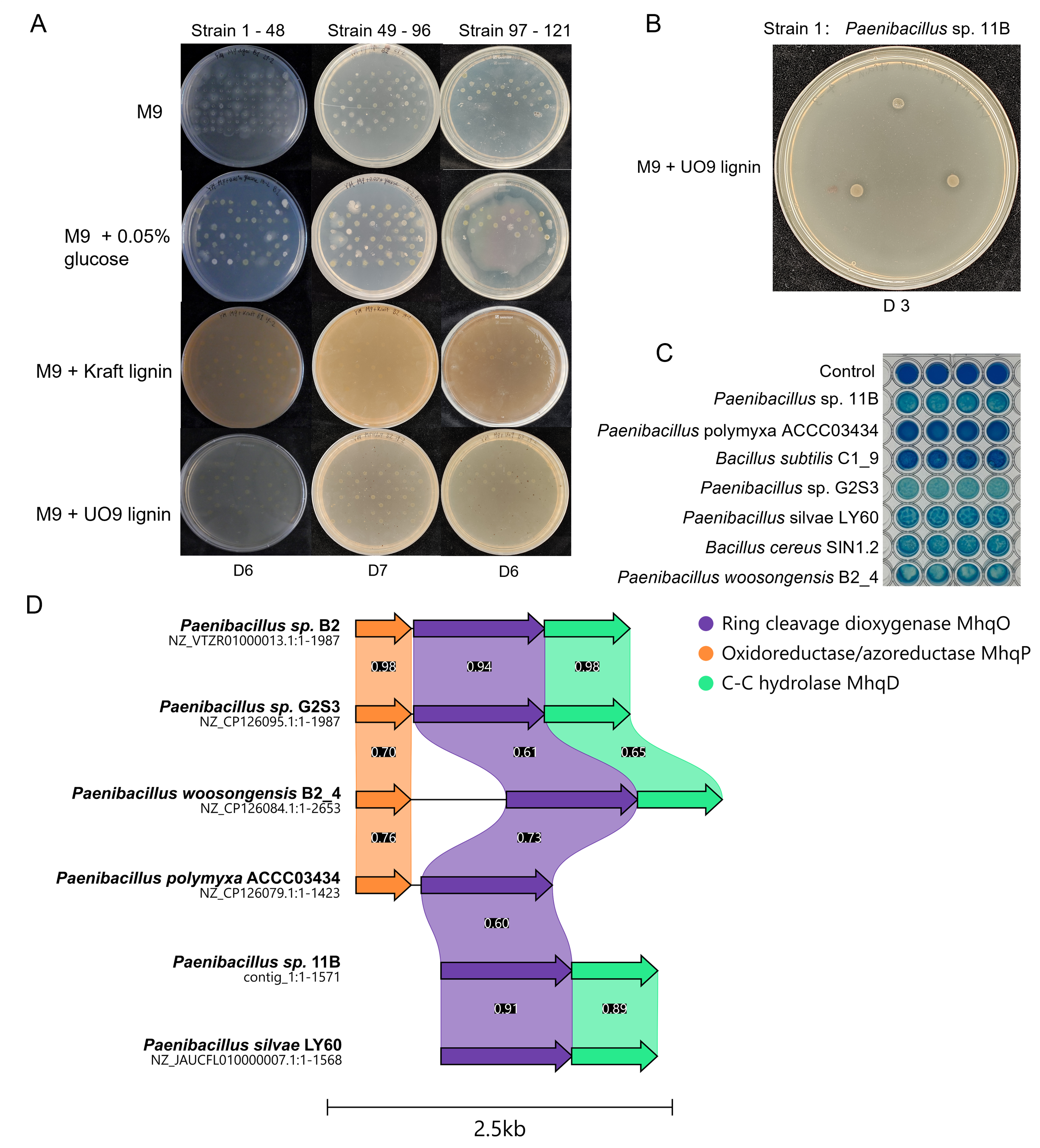
Bottom-up screening of *Bacillaceae* strains for lignin-associated growth, ligninolytic activity, and genetic potential. (A) Growth screening of *Bacillales* strain collection stamped onto M9 agar medium supplemented with different carbon sources, including no added carbon source (M9), 0.05% (w/v) glucose, kraft lignin, or U09 beechwood lignin (from top to bottom rows). (B) Representative clear halo formation by *Paenibacillus* sp. 11B on M9 agar medium supplemented with U09 lignin after 3 days of incubation. (C) Qualitative decolorization of the lignin-like dye methylene blue (MB) by the selected *Bacillales* strains. Wells containing dye without bacteria served as controls. (D) Comparative genomic organization of the *mhq* gene cluster in selected *Paenibacillus* strains. The numbers denote protein similarity values.

To further investigate the ligninolytic potential of each strain independently of lignin utilization, decolorization of synthetic lignin-mimicking dye methylene blue was tested (MB, Figure 2C) (Husain, 2006). Including strain 11B, all *Paenibacillus* strains from the collection, together with two randomly selected strains were tested in a rapid MB decolorization assay with a short incubation time (20 min). Notably, *Paenibacillus* isolates displayed very fast MB decolorization or absorption, with strain G2S3 showing the fastest reduction within 20 min.

As MB is not a direct proxy for lignin degradation, we next explored the presence of genes potentially involved in ligninolytic activity of these *Paenibacillus* isolates. In particular, pathways involved in the cleavage of lignin-derived aromatic structures could provide a direct link to lignin catabolism. A recent study by Fanitsios et al. reported that the lignin-degrading strains *Paenibacillus* sp. B2, *Agrobacterium* sp. B1, and *Ochrobactrum* sp. each carry the *mhqO* gene encoding a putative ring-cleaving dioxygenase (Fanitsios et al., 2025). These strains were shown to degrade the biphenyl-containing lignin fragment 5,5′-di (dehydrovanillic acid) (DDVA) on solid media. To investigate whether similar catabolic potential exists in our *Paenibacillus* isolates, we retrieved the protein sequences encoded within the *mhq* gene cluster from *Paenibacillus* sp. B2, including WP_149645412 (encoding C–C hydrolase MhqD), WP_149645413 (encoding ring-cleaving dioxygenase MhqO), and WP_149645414 (encoding oxidoreductase/azoreductase MhqP), and compared with five *Paenibacillus* strains in our collection. This comparative analysis revealed that the proteins encoded in the gene cluster, particularly the ring-cleavage dioxygenase MhqO, are highly conserved within the *Paenibacillus* genus (Figure 2D), indicating that the genetic potential for this aromatic catabolic pathway is widespread in the genus

### Top-down selection of lignin-enriched biofilm communities

The biofilm-forming ability and ligninolytic enzyme repertoire of *Bacillales* are both likely to contribute to effective lignin biodegradation. Consequently, only using a bottom-up selection strategy may overlook strains that do not exhibit strong ligninolytic activity in isolation but nonetheless play an important ecological role in a community context. To overcome this limitation, we complemented the bottom-up screening with a top-down selection approach (Figure 1). We established enrichment cultures with and without inoculation of the 121-strain *Bacillales* library, and with or without lignin supplementation. After 7 days of incubation, flasks inoculated with the *Bacillales* library developed a ring-shaped biofilm along the inner wall of the flasks (Supplementary Fig. S1). We hypothesize that biofilm-forming strains facilitate lignin biodegradation by retaining lignin particles within the matrix and enabling localized enzymatic activities.

The biofilm ring was harvested for both single-colony isolation and community DNA extraction. Sanger sequencing of cultured isolates identified several recurrent taxa, including *Lysinibacillus pakistanensis* LY18, *Bacillus subtilis* D9_B_56, *Bacillus bombysepticus* ACCC04323, *Bacillus sonorensis* YX13, *Bacillus velezensis* MB7_B13, and *Bacillus licheniformis* D9_B_45. Nanopore sequencing provided a complementary community-level perspective across five biological replicates, *Bacillus paralicheniformis* CEW_1W consistently dominated the enriched biofilms (Figure 3). Importantly, *Paenibacillus* sp. 11B, selected in the bottom-up assay for its high ligninolytic potential, was the second most abundant species in the sequencing-based analysis across all enriched biofilm samples, underscoring its likely contribution to lignin degradation. Integrating evidence from both cultivation-based identification and metataxonomic profiling, we selected 15 strains as candidates for SynCom assembly (Table 1).

**Figure 3.**
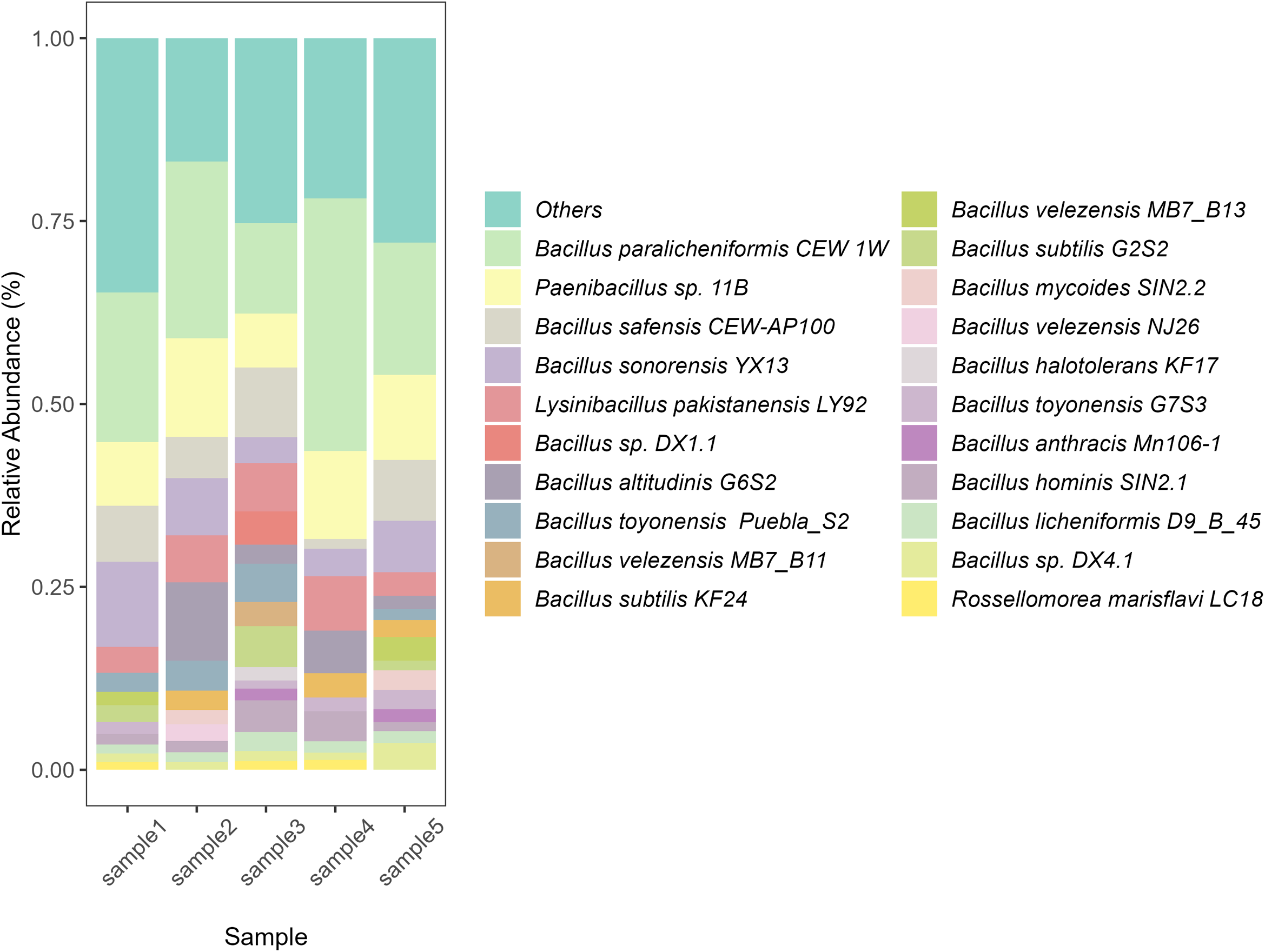
Relative abundance of taxa in the ring-shaped *Bacillales* biofilm community. Relative abundance lower than 1% was classified as ‘others’, Each bar represents an individual biological replicate (n=5).

**Table 1.** Summary of candidate strains for lignin degradation SynCom.

| NO. | TAXONOMY | STRAIN | ASSEMBLY | ISOLATION SOURCE |
| --- | --- | --- | --- | --- |
| 1 | *Paenibacillus* sp. | 11B | GCA_030487755.1 | head of a worker termite |
| 2 | *Bacillus bombysepticus* | ACCC04323 | GCA_030348965.1 | soil |
| 3 | *Lysinibacillus pakistanensis* | LY18 | GCA_030348485.1 | cucumber rhizosphere |
| 4 | *Bacillus subtilis* | D9_B_56 | GCA_030348595.1 | grassland soil |
| 5 | *Bacillus sonorensis* | YX13 | GCA_031324045.1 | rhizosphere soil |
| 6 | *Bacillus licheniformis* | D9_B_45 | GCA_030348625.1 | grassland soil |
| 7 | *Bacillus velezensis* | MB7_B13 | GCA_030348465.1 | grassland soil attached to a mushroom |
| 8 | *Bacillus paralicheniformis* | CEW_1W | GCA_030123025.1 | marine or sediment |
| 9 | *Bacillus safensis* | CEW_AP10<br>0 | GCA_030122965.1 | marine or sediment |
| 10 | *Lysinibacillus pakistanensis* | LY92 | GCA_030123265.1 | cucumber rhizosphere |
| 11 | *Bacillus* sp. | DX1.1 | GCA_030348225.1 | field soil |
| 12 | *Bacillus altitudinis* | G6S2 | GCA_030123125.1 | pine forest soil |
| 13 | *Bacillus toyonensis* | Puebla_S2 | GCA_030167055.1 | semidesertic area with dry sandy soils, around 15 cm from the trunk of a lemon tree (5 cm deep) |
| 14 | *Bacillus velezensis* | MB7_B11 | GCA_030123285.1 | grassland soil attached to a mushroom |
| 15 | *Bacillus subtilis* | KF24 | GCA_030123145.1 | rhizosphere soil |

### Biofilm formation and lignin-like dye decolorization–guided selection of *Bacillales* strains for the SynCom

A total of 15 strains were primarily selected motivated by bottom-up and top-down selection results (Table 1). As the community member size increases, the number of interactions within a SynCom arises non-linearly, reducing the tractability and predictability of community-level functions. To reduce the community complexity, single-strain assays for biofilm formation and lignin-like dye decolorization were performed (Fig. S2).

Based on crystal violet staining, 2 strains (D9_B_45 and G6S2) showed strong biofilm formation ability, 3 strains (CEW_1W, Puebla_S2, and KF24) showed moderate biofilm formation, whereas the remaining strains had only weak or negligible biofilm formation (Fig. S2A). To investigate ligninolytic potential independently of lignin utilization, decolorization of synthetic lignin-like dyes, methylene blue (MB) and Azure B (AB) was monitored for the 15 *Bacillales* strains (Fig. S2B-C). MB and AB are used as proxy assays and are not interpreted as direct evidence. Most strains efficiently decolorized MB, whereas strain YX13 showed only very limited decolorization. In contrast, overall decolorization of AB was lower than that of MB, but strain Puebla_S2 still achieved a relatively high decolorization efficiency (close to 80%), while strain YX13 again performed poorly. In addition, strains ACCC04323 and LY18 showed negligible biofilm formation and only moderate dye decolorization, while strain DX1.1 exhibited very slow growth. These strains were therefore not retained in the final SynCom. Taken together, strains ACCC04323, LY18, YX13 and DX1.1 showed poor biofilm formation and/or low dye decolorization and were therefore excluded from the final SynCom design.

Based on the results, a total 11 strains were shown to produce biofilm or decolorize lignin-like dyes were selected as a SynCom for further analysis (Figure 4A).

**Figure 4.**
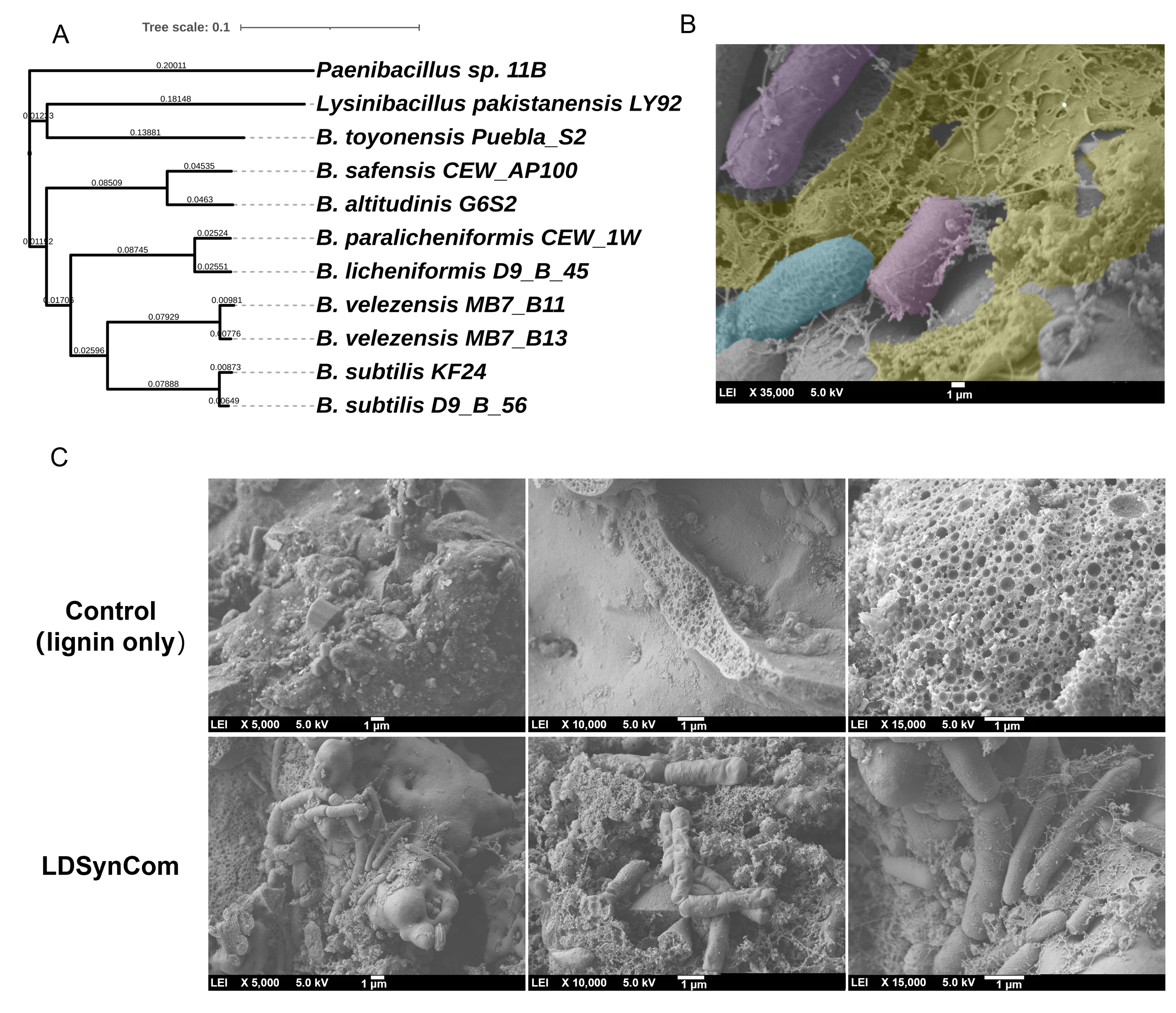
Composition and biofilm formation of the LDSynCom. (A) Phylogenetic tree of the 11 *Bacillales* strains selected for LDSynCom assembly based on combined bottom-up and top-down selection strategies. The tree was constructed using assembled complete genomes under NCBI Bioproject PRJNA960711; branch lengths indicate sequence divergence. (B) Representative pseudo-colored SEM micrograph. The yellow-colored regions indicate the extracellular biofilm matrix, while different colors highlight bacterial cells with distinct morphologies corresponding to different LDSynCom members. Colors were added for visualization purposes only. Scale bar denotes 1μm. (C) SEM images of lignin particles incubated under sterile conditions (abiotic control, lignin only, top row) or in the presence of the LDSynCom after 10 days of incubation (bottom row). Representative images at increasing magnifications are shown from left to right, respectively. Scale bars denote 1μm.

### Direct visualization of mixed species biofilm formation on lignin

Scanning electron microscopy (SEM) was used to confirm that the lignin degrading SynCom (LDSynCom) can produce biofilms on lignin surfaces (Figure 4B and C). After 10 days of incubation in lignin-supplemented M9 medium, the characteristic ring-shaped biofilm formed along the inner wall of the shake flasks was collected and used for SEM analysis. In the non-inoculated control, the complex structure of the lignin matrix was observed solely (Figure 4C). In the LDSynCom-inoculated samples, well-developed biofilm structures could be observed attached to the surface of lignin particles. Within these structures, bacterial cells were embedded in an extracellular matrix-like network, indicating stable surface-associated growth. Notably, bacterial cells with distinct morphologies could be distinguished, consistent with the multispecies composition of the LDSynCom. The observation of biofilm structures on lignin particles indicates that the LDSynCom contains strong biofilm formers that generate EPS matrix, potentially functioning as a localized “digestive system” that enhances close cell–substrate contact.

### *Paenibacillus* sp. 11B contributes to phenolic release from lignin in LDSynCom

We hypothesized that within the LDSynCom, distinct strains play complementary functional roles. Some members act as strong biofilm formers. Other strains function as “degraders” that secrete ligninolytic enzymes, exemplified by *Paenibacillus* sp. 11B, which shows pronounced degradation activity on lignin-containing agar. Consistent with these potential cooperative interactions, *Paenibacillus* sp. 11B exhibited little biofilm-forming capacity in both crystal violet staining and shake-flask assays (Fig. S3), whereas strains such as *B. licheniformis* D9_B_45 formed robust ring-shaped biofilms but moderate degradation ability (Fig. S3).

To test whether strain 11B contributes to lignin-associated activity within the consortium, we performed a drop-out experiment in which this strain was removed from the LDSynCom and assessed the resulting impact on lignin degradation potential (Figure 5). LDSynCom, LDSynCom without 11B (LDSynCom-11B), and 11B alone were inoculated in M9 medium supplemented with 0.2% glucose and 0.1% lignin. Uninoculated medium was used as the control. Under these conditions, LDSynCom showed sustained growth and reached the highest final biomass among all cultures over the 41-days incubation (Figure 5B). Removing strain 11B from the LDSynCom reduced the overall growth, whereas strain 11B alone showed limited biomass accumulation compared with the full LDSynCom (Figure 5B).

**Figure 5.**
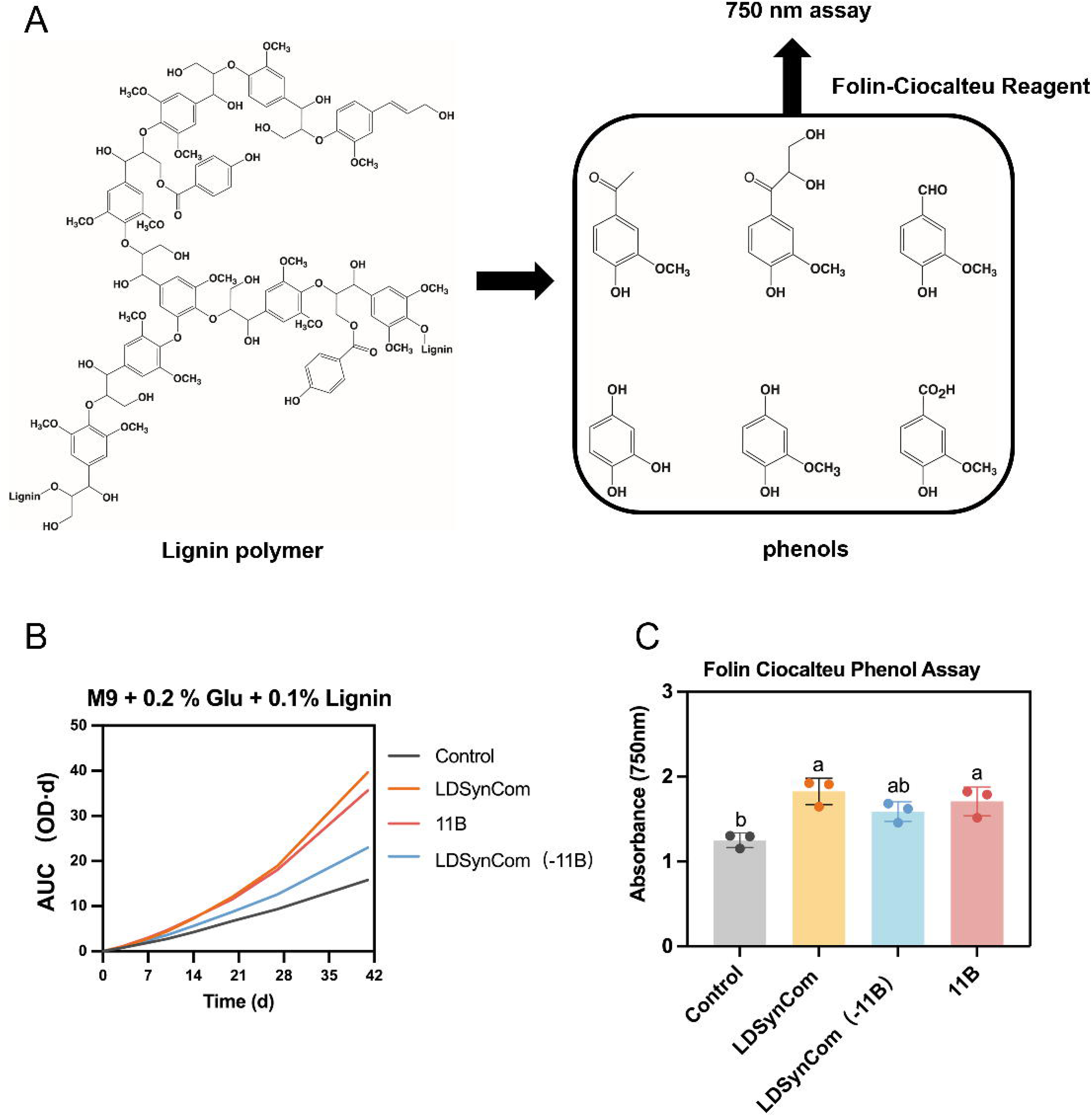
Batch culture assay of LDSynCom-mediated lignin depolymerization. (A) Scheme of assay for small molecule phenolic products from polymeric lignin. (B) Area under the growth curve (AUC) of LDSynCom, LDSynCom-11B (LDSynCom without strain 11B), and strain 11B alone grown in M9 medium supplemented with 0.2% (w/v) glucose and 0.1% (w/v) lignin. AUC values were calculated from OD□□□ measurements over the 42-day incubation period. (C) Total phenols released at day 28, quantified by the Folin–Ciocalteu assay (FCA); uninoculated medium served as the control. Data in (B) and (C) are shown as mean ± SD of n = 3 biological replicates. In (C), statistical differences among groups were assessed by one-way ANOVA with Tukey–Kramer’s post hoc test; groups sharing the same letter are not significantly different, whereas different letters indicate significant differences (P < 0.05).

Depolymerization of lignin can result in the release of soluble phenolic compounds derived from its complex aromatic structure (Rashid and Bugg, 2021). To further verify possible release of phenolic compounds from lignin during cultivation, FCA was applied to quantify total released phenols signals at A750 (Figure 5A). The LDSynCom samples produced significantly higher phenolic signals than the uninoculated control and it was higher than LDSynCom-11B and 11B samples (Figure 5C). Thus, LDSynCom promotes greater phenolic accumulation, and the presence of strain 11B contributes to this activity of the consortium. Compared with *Paenibacillus* sp. 11B grown alone, the LDSynCom achieved a higher biomass yield, further supporting the functional complementarity within the community. While strain 11B provides the core lignin-associated activity through enzyme secretion, its performance seems to be enhanced in the presence of biofilm formers.

### Proteomics supports member-dependent contributions to lignin depolymerization

To functionally map lignin-associated activities within the LDSynCom, we performed community-level proteomics comparing lignin-supplemented (M9 + 0.2% glucose + lignin) and control conditions (M9 + 0.2% glucose). In both treatments, 0.2% glucose served as the primary readily available carbon source to support initial bacterial growth. We conducted differential expression analysis on 3,986 protein groups following removal of 24 contaminant proteins (Figure 6A). Among these, 1,171 protein groups were exclusively detected under lignin-supplemented conditions, whereas 1,619 were unique to the control condition (Dataset S2). Representative examples of lignin-specific proteins include an FMN-dependent NADH azoreductase AzoRB (WP_003228400.1) in strain KF24 and a LolA family protein (WP_283872006.1) in strain LY92, both exclusively detected under lignin-supplemented conditions.

**Figure 6.**
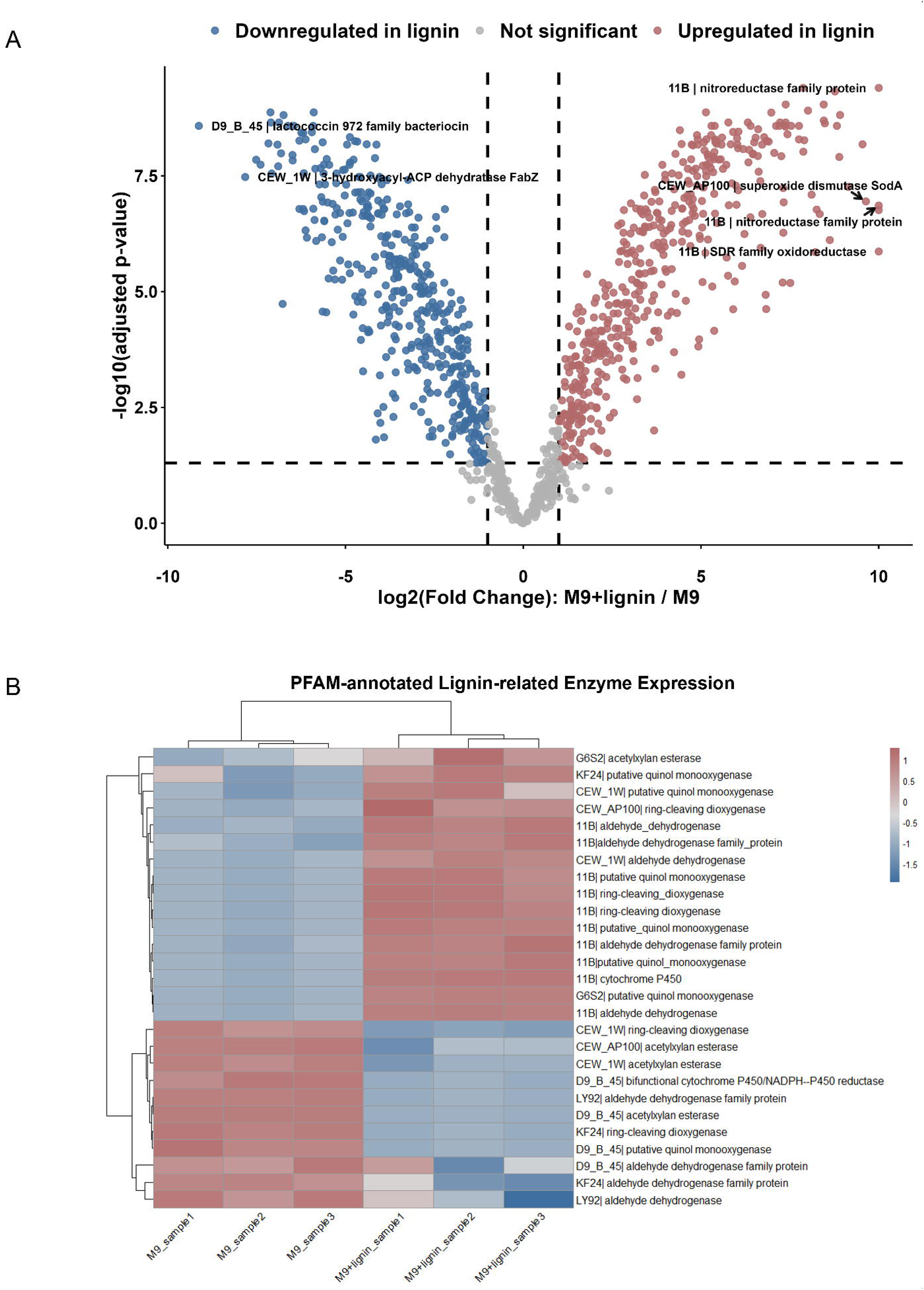
Lignin-induced functional reprogramming and ligninolytic enzyme expression shifts in the LDSynCom. (A) Volcano plot showing the global proteomic response to lignin supplementation. Log2 fold change of log-transformed protein abundance (M9 + lignin vs. M9) is plotted against −log10(padj) for protein groups detected in both conditions. Significantly differentially abundant proteins are highlighted in red (upregulated in lignin) and blue (downregulated in lignin), while non-significant proteins are shown in grey. The most strongly regulated proteins are annotated. (B) Heatmap representation of lignin-degrading enzyme expression under M9 and M9+lignin conditions. Z-score normalization was applied to eliminate baseline abundance differences across proteins, and hierarchical clustering was used to identify co-regulated enzyme clusters.

The remaining 1,196 protein groups were shared between both conditions. Of the shared proteins, 900 (75.2%) exhibited significant differential abundance (p < 0.05), including 459 proteins with higher abundance in the presence of lignin and 441 proteins showing higher abundance in the M9 control. Lignin-associated differences were observed within LDSynCom. Lactococcin (WP_003177941.1) in D9_B_45 and 3-hydroxyacyl-ACP dehydratase FabZ (WP_003185902.1) in CEW_1W showed significantly higher abundance under M9 control conditions. In contrast, a nitroreductase family protein and an SDR family oxidoreductase showed significantly higher abundance in strain 11B under lignin-supplemented conditions (WP_301881884.1, WP_301879059.1, WP_079349695.1). Similarly, in CEW_AP100, superoxide dismutase SodA (WP_007501264.1) exhibited increased abundance in the presence of lignin. The high proportion of differentially abundant proteins reflects the profound impact of lignin supplementation on the LDSynCom proteome.

To further characterize functional responses to lignin supplementation, differential expression analysis was performed on 3,986 protein groups. The top 50 differentially expressed proteins were visualized in a hierarchically clustered heatmap (Fig. S4). In addition, 27 PFAM-annotated lignin degradation-associated enzymes were identified across conditions (Figure 6B). Most lignin-associated enzymes displayed increased abundance under lignin-supplemented conditions. Notably, a substantial fraction of the lignin-induced proteins originated from strain 11B, including ring-cleaving dioxygenases that are conserved among *Paenibacillus* species and commonly identified in previously characterized lignin-degrading bacteria (Fanitsios et al., 2025). This abundance response indicates that the *mhqO* homolog encoded by strain 11B is induced in the presence of lignin and likely contributes to aromatic ring cleavage during lignin depolymerization. Moreover, putative quinol monooxygenases and aldehyde dehydrogenases were upregulated across multiple strains, including strain 11B, CEW_1W, and KF24.

## Discussion

Synthetic microbial communities have been extensively investigated and discussed in gut microbiota and plant microbiome research, but until now not explored in detail in lignin degradation. To our knowledge, this study represents one of the first systematic attempts to deploy a rationally designed SynCom in lignin degradation. Building upon established SynCom design principles from other fields, we integrated top-down and bottom-up strategies for the construction of SynCom. This dual framework proved critical for capturing emergent community-level properties that would likely be overlooked by either approach alone.

The top-down strategy enriched for strains exhibiting strong biofilm formation capacity, a trait that might be central to lignin degradation. Increasing evidence suggests that lignin and lignocellulose degradation in natural and engineered systems is closely linked to biofilm-associated lifestyles rather than planktonic growth. Previous work has shown that biofilm formation on lignin matrices correlates with elevated metabolic activity, including increased oxygen consumption and carbon dioxide production, and may enhance enzyme–substrate interactions and tolerance to toxic intermediates generated during lignocellulose processing (Meslé et al., 2022). More broadly, studies on plastic degradation have demonstrated that biofilms concentrate extracellular enzymes at the substrate interface, thereby improving degradation efficiency (Howard and McCarthy, 2023). Further experiments will be required to test whether lignin-degrading enzymes are similarly enriched within the biofilm matrix, including biofilm-disruption assays, EPS fractionation, and comparison of enzyme localization and activity across the matrix. Beyond validating this mechanism, biofilm formation also offers a practical handle for improving degradation performance. Experimental evolution has been used as a selection strategy to develop biofilm-based SynComs with enhanced plastic-degrading activity(S. Li et al., 2025). Therefore, we hypothesize that iterative selection of a biofilm-forming SynCom on lignin matrices through experimental evolution will promote the emergence of a stable community with increased biofilm-forming capacity thereby enhancing substrate colonization, enzyme–substrate proximity, and metabolic cooperation, ultimately leading to improved lignin degradation.

The bottom-up selection strategy identified *Paenibacillus* as a genus of interest for lignin-related functions, with methylene blue decolorization used as a rapid, visual, and scalable primary filter for prioritizing SynCom candidates from a collection of 121 strains. Confirming lignin-degrading function in individual strains will require direct enzymatic assays and quantification of lignin-derived degradation products. Comparative genomic analysis revealed a conserved *mhqPOD* gene cluster within multiple *Paenibacillus* strains, indicating a genetically encoded capacity for processing lignin-derived aromatic compounds. Consequently, Fanitsios et al. demonstrated by quantitative PCR that an operon comprising five *mhq* genes in *Paenibacillus sp.* B2 was strongly induced with 400-1000 fold overexpression in the presence of lignin-derived compound 5,5’-di (dehydrovanillic acid) (Fanitsios et al., 2025). Such pronounced transcriptional response underscores the potential relevance of *mhq* operon involved in lignin degradation. Consistent with the transcriptional patterns, our proteomic analysis further revealed increased abundance of lignin-associated enzymes under lignin-supplemented conditions, including a homolog of the ring-cleaving dioxygenase MhqO encoded by strain 11B. The concordant transcriptional and proteomic responses provide partial functional support that the *mhq* cluster is actively involved in the cellular response to lignin and may contribute to aromatic compound breakdown during lignin depolymerization. Future work should focus on targeted gene knockouts or functional perturbation of the *mhqPOD* cluster to establish causal links between this operon and lignin-associated metabolic activity. Notably, strain 11B was isolated from the head of a worker termite (Song et al., 2024), an ecological niche renowned for lignocellulose turnover. The ecological origin of this strain may therefore reflect evolutionary adaptation to lignin-rich environment (Kudo, 2009). Exploring termite associated microbiomes might provide a rich reservoir of lignin-degrading specialists and inform the rational design of next-generation lignin degradation SynComs.

In this study, a targeted drop-out experiment focusing on strain 11B, together with community-level proteomic profiling, supported its contribution to lignin degradation within the consortium. While strain 11B may represent a primary ligninolytic “degrader” within the LDSynCom, lignin-like dye decolorization assays indicate that multiple community members possess degradative potential. This suggests that functional roles within the consortium are not strictly partitioned, and some strains may simultaneously contribute to biofilm formation and lignin transformation. Future drop-out and add-in experiments on different SynCom members would help resolve the functional role of each community member and deepen our mechanistic understanding of community-level lignin metabolism. In addition, genome-scale metabolic models could be applied to predict the metabolic interactions between SynCom members and to identify optimal strain combinations. Combining these approaches with time-resolved community profiling, such as amplicon sequencing at successive cultivation time points, would further capture the temporal dynamics and stability of the community during lignin degradation. Together, such analyses could identify metabolic cross-feeding interactions and clarify the extent of functional redundancy within the consortium. Ultimately, this knowledge could guide rational design of a minimal yet high-performing and stable ligninolytic SynCom.

To assess lignin degradation, we applied two rapid spectrophotometric assays, UV absorbance at 280 nm and the Folin-Ciocalteu assay (Z. Xu et al., 2018b). Lignin exhibits a strong UV band around 280 nm that originates primarily from aromatic rings and phenolic groups (J. Li et al., 2025). Consequently, absorbance at 280 nm (A280) is widely used as a rapid measurement for lignin degradation (Z. Xu et al., 2018b). However, in our work A280 increased over time (Fig. S5). This increase is still biologically plausible, because lignin degradation could release low-molecular-weight aromatic fragments into the medium, raising A280. Moreover, A280 is not lignin-specific, as proteins, nucleic acids, secreted enzymes, and cell lysis can all contribute to absorbance in this region. Meanwhile, lignin is depolymerized to release aromatic rings and phenolic compounds which further increase A280. It should be noted that both A280 and the Folin–Ciocalteu assays provide indirect measures of lignin-associated changes and do not provide direct structural evidence of lignin depolymerization. Further characterization using techniques such as LC–MS/GC–MS (Liquid Chromatography–Mass Spectrometry/Gas Chromatography–Mass Spectrometry), GPC/SEC (Gel Permeation Chromatography/Size Exclusion Chromatography), and 2D-HSQC NMR (Two-Dimensional Heteronuclear Single Quantum Coherence Nuclear Magnetic Resonance) would be required to identify lignin-derived products and characterize structural changes in lignin. These complementary analyses will be important for more direct assessment of lignin depolymerization in future studies.

Proteomic analysis further enlightened the response of SynCom to lignin supplementation. Among the differentially abundant proteins, FMN-dependent NADH azoreductase (AzoRB) in *B. subtilis* KF24 was exclusively detected under lignin-supplemented conditions, suggesting its abundance is strongly associated with lignin supplementation. Salvachúa *et al*. reported enrichment of an FMN-dependent NADH azoreductase in outer membrane vesicles (OMVs) of *Pseudomonas putida* KT2440 cultivated in lignin-containing medium (Salvachúa et al., 2020), suggesting that lignin-containing conditions can favor the enrichment of reductases. FMN-dependent NADH azoreductase is also known as a member of azoreductases (Matsumoto et al., 2010), which plays a central role in metabolism of azo dyes. Moreover, azoreductases can be potentially involved in the modification of both oligomeric and monomeric aromatic or phenolic compounds (Gonçalves et al., 2013; Matsumoto et al., 2010). Taken together, the enrichment of AzoRb in KF24 suggests that this is a putative candidate involved in downstream metabolism of lignin-derived aromatics released. Strain 11B also showed increased abundance of multiple putative ring-cleavage dioxygenases under lignin supplementation. Ring-cleavage dioxygenases catalyze the opening of aromatic rings, a critical step that channels aromatic compounds into central metabolism (Johnson and Beckham, 2015; Semana and Powlowski, 2019). Their enrichment in 11B is consistent with its role in downstream catabolism of lignin-derived aromatic compounds. Importantly, the lignin-related enzymes identified in strain 11B provide a set of candidate enzymes for future elucidating the mechanism and pathways of lignin degradation.

## Concluding remarks

Here, we provide the first analysis of both the ligninolytic enzymatic potential and biofilm-forming capacity across a diverse collection of *Bacillales*. We introduce an integrated top-down and bottom-up screening framework that enables the rational discovery and assembly of lignin-degrading SynCom. We hypothesize that LDSynCom forms lignin-associated biofilms, and that biofilm formation is functionally linked to lignin degradation. Consistent with this hypothesis, the LDSynCom formed biofilm structures on the surface of complex lignin structure, indicating close physical association between cells and lignin particles. Importantly, Folin–Ciocalteu assays confirmed increased accumulation of phenolic products in the presence of LDSynCom. Proteomic profiling identified proteins annotated as lignin-related enzymes in the LDSynCom and revealed lignin-associated differences in protein abundance, including a higher abundance of ring-cleavage dioxygenases in 11B. Together, our results establish an ecologically informed strategy for constructing functionally partitioned SynComs, offering a powerful platform for mechanistic studies of lignin breakdown and for the development of future lignin valorization technologies.

## Acknowledgements

Q.Z. was funded by China Scholarship Council. X.Xin was funded by European Union via ERC Advanced Grant 101055020-COMMUNITY to G.P.v.W. The position of X.Xu was funded by a Startup from Institute of Biology to Á.T.K. The laboratory of Á.T.K. is funded by the European Union (ERC, MicroClock, 101166968). Views and opinions expressed are, however, those of the author(s) only and do not necessarily reflect those of the European Union or the European Research Council Executive Agency. Neither the European Union nor the granting authority can be held responsible for them. We thank dr. André van Zomeren, TNO, The Netherlands, for supplying beechwood lignin samples.

---

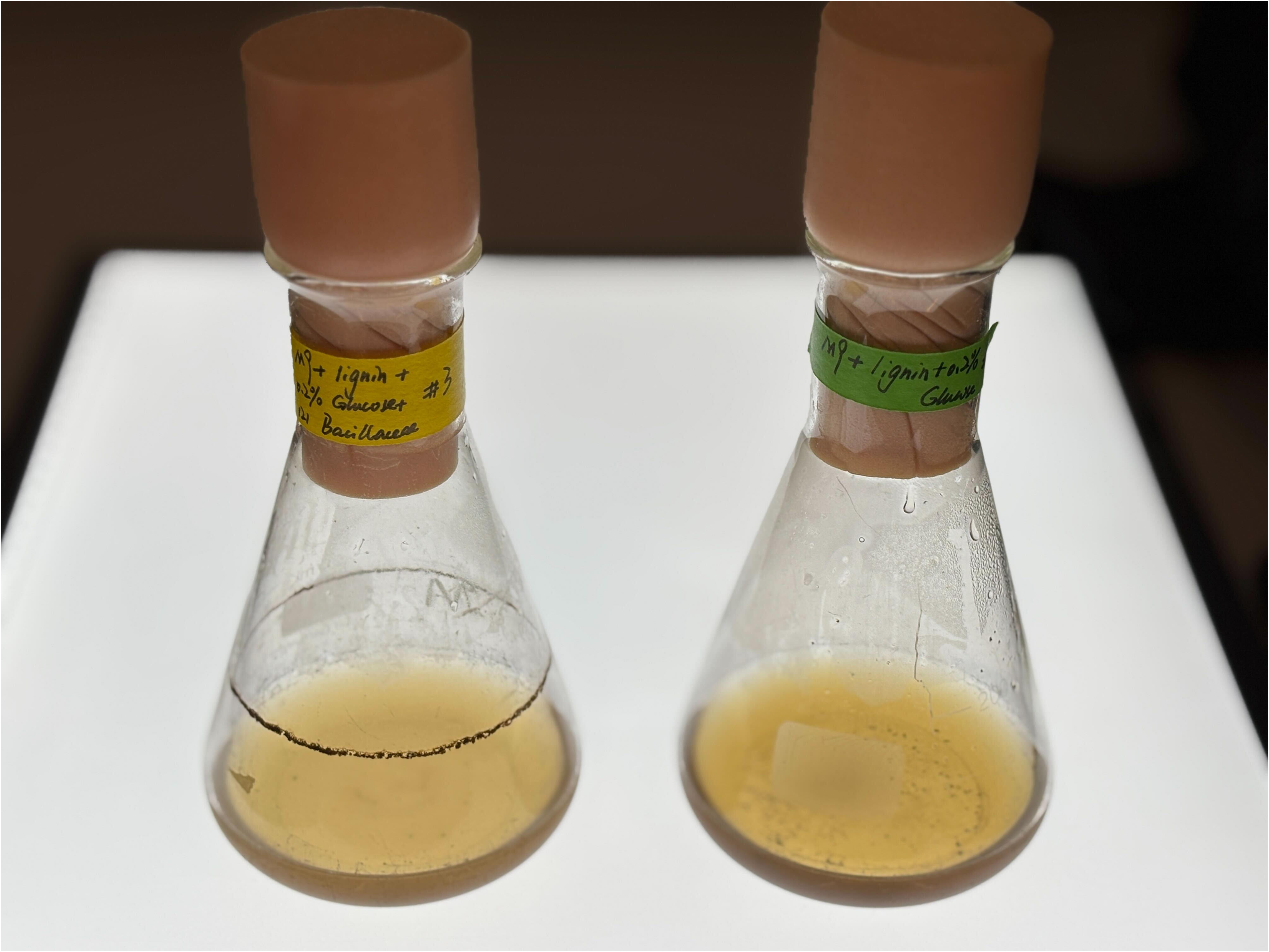

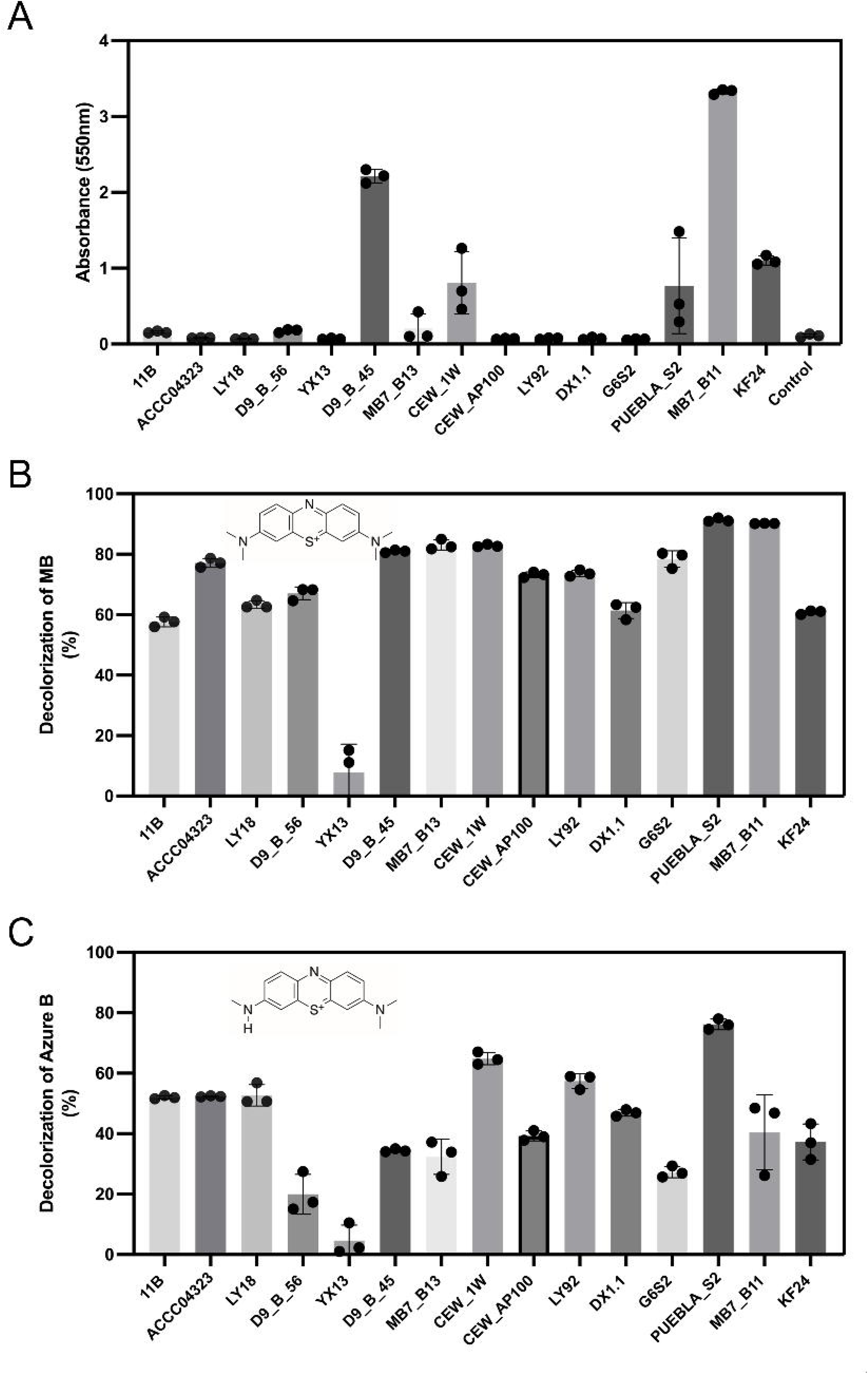

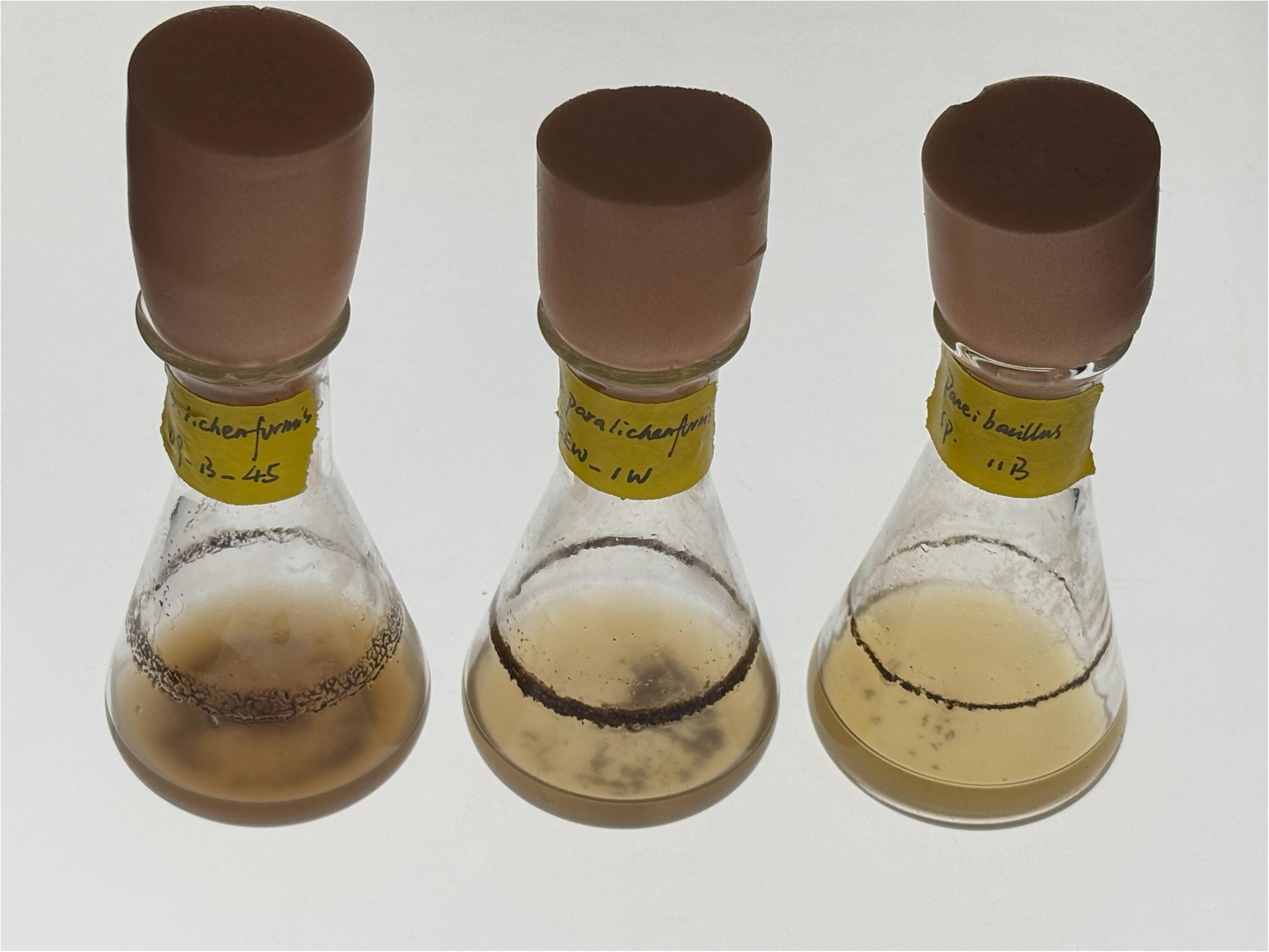

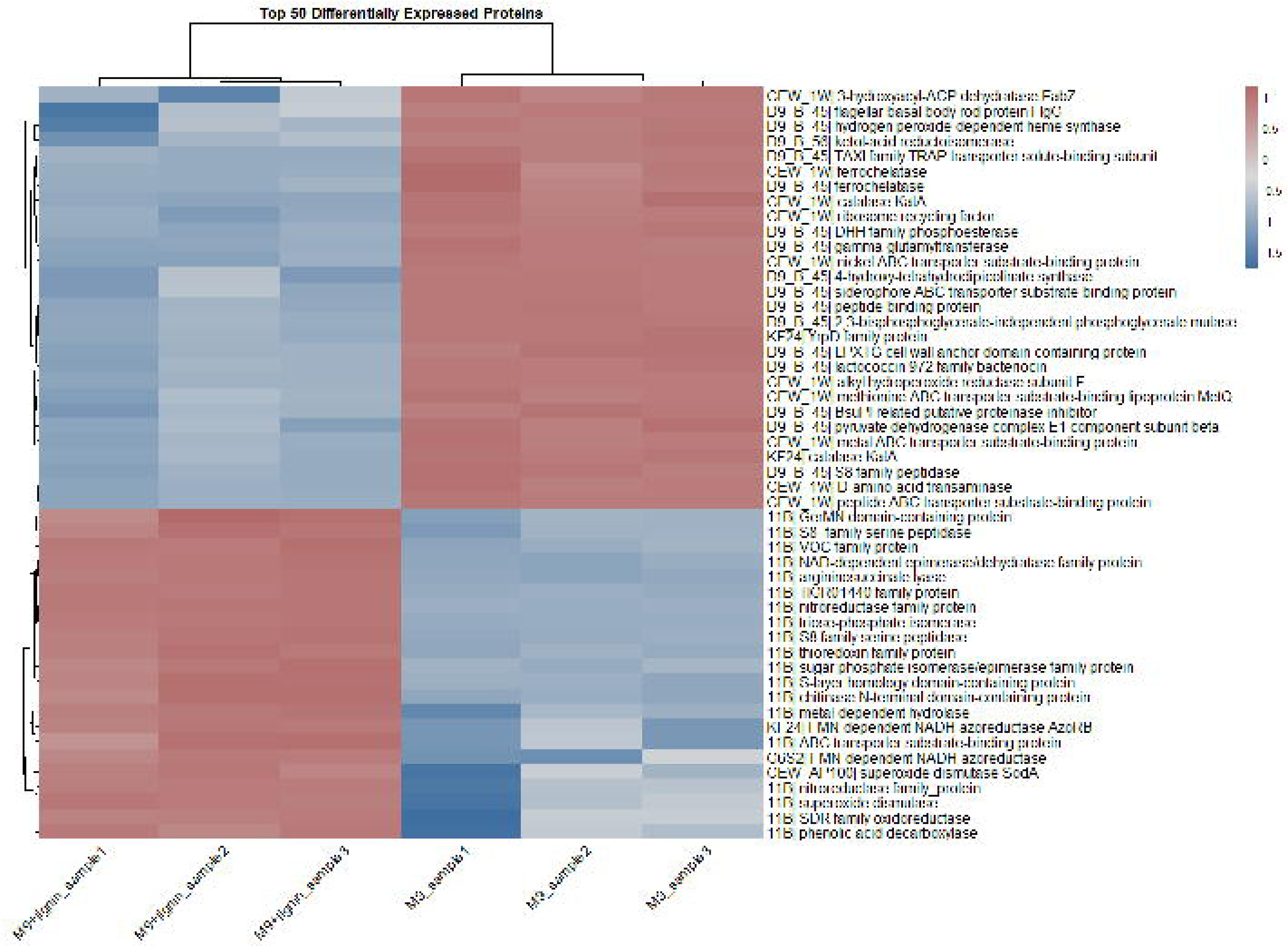

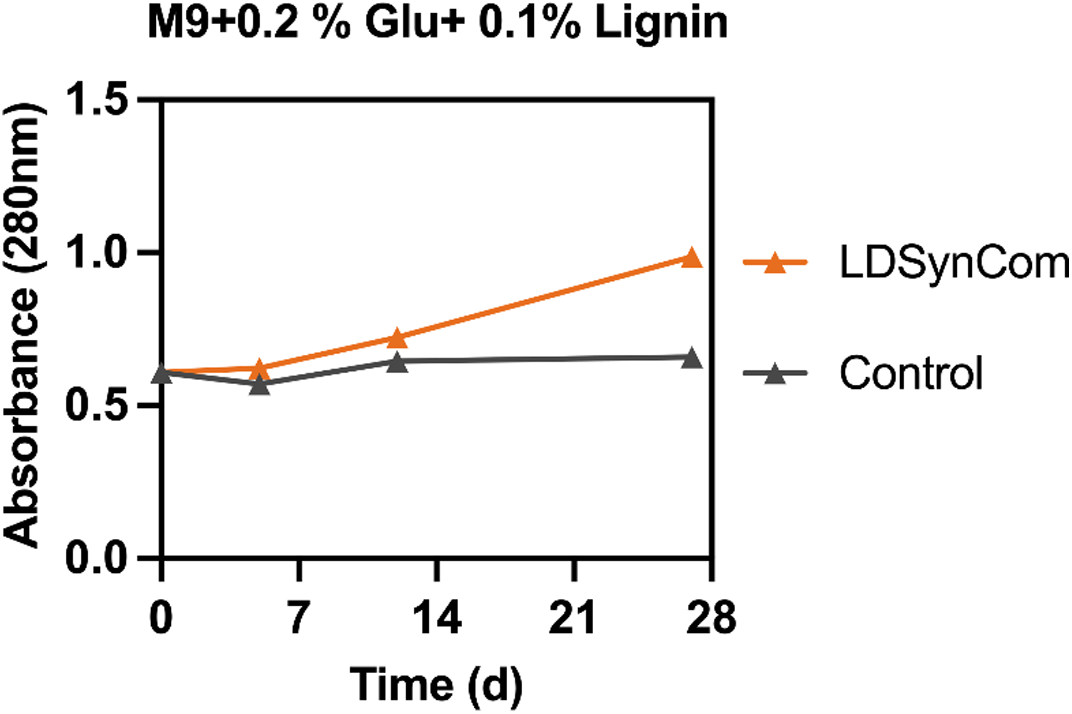

