## Supplementary File for "Ecologically inspired assembly of a *Bacillales* synthetic community for lignin degradation"

Table S1 The 121 *Bacillales* strains collection.

| 1 | *Paenibacillus* sp. | (11B) AKRM2 |
| --- | --- | --- |
| 2 | *Bacillus bombysepticus* | ACCC01970 |
| 3 | *Halobacillus* sp*.* | ACCC02827 |
| 4 | *Virgibacillus halodenitrificans* | ACCC02857 |
| 5 | *Brevibacillus parabrevis* | ACCC02960 |
| 6 | *Brevibacillus agri* | ACCC03016 |
| 7 | *Paenibacillus polymyxa* | ACCC03434 |
| 8 | *Lysinibacillus sphaericus* | ACCC03577 |
| 9 | *Bacillus bombysepticus* | ACCC04323 |
| 10 | *Peribacillus* sp*.* | ACCC06369 |
| 11 | *Ureibacillus composti* | AQ11 |
| 12 | *Mesobacillus* sp. | AQ2 |
| 13 | *Bacillus subtilis* | b15 |
| 14 | *Fictibacillus enclensis* | b19 |
| 15 | *Bacillus pumilus* | B2_10 |
| 16 | *Paenibacillus woosongensis* | B2_4 |
| 17 | *Fictibacillus* sp. | b24 |
| 18 | *Bacillus subtilis* | b4 |
| 19 | *Bacillus subtilis* | C1_13 |
| 20 | *Bacillus subtilis* | C1_9 |
| 21 | *Bacillus velezensis* | Canada_S1 |
| 22 | *Priestia flexa* | (CCW CN82) CU15 |
| 23 | *Bacillus altitudinis* | (CES-OCA-19) CU10 |
| 24 | *Bacillus paralicheniformis* | (CEW 1W) CU9 |
| 25 | *Bacillus safensis* | (CEW AP102) CU8 |
| 26 | *Bacillus safensis* | (CEW AP100) CU11 |
| 27 | *Neobacillus* sp. | CF12 |
| 28 | *Peribacillus frigoritolerans* | CF13 |
| 29 | *Peribacillus frigoritolerans* | CF15 |
| 30 | *Peribacillus frigoritolerans* | CF20 |
| 31 | *Peribacillus frigoritolerans* | CF29 |
| 32 | *Bacillus bombysepticus* | Cuernavaca_S2 |
| 33 | *Bacillus toyonensis* | Cuernavaca_S4 |
| 34 | *Bacillus cereus* | D5_B_69 |
| 35 | *Bacillus pumilus* | D8_B_31 |
| 36 | *Peribacillus simplex* | D8_B_37 |
| 37 | *Peribacillus simplex* | D8_B_41 |
| 38 | *Bacillus cereus* | D8_B_42 |
| 39 | *Bacillus mycoides* | D8_B_46 |
| 40 | *Bacillus cereus* | D8_B_47 |
| 41 | *Bacillus wiedmannii* | D9_B_34 |
| 42 | *Bacillus subtilis* | D9_B_36 |
| 43 | *Bacillus licheniformis* | D9_B_45 |
| 44 | *Bacillus altitudinis* | D9_B_49 |
| 45 | *Bacillus subtilis* | D9_B_56 |
| 46 | *Bacillus subtilis* | D9_B_66 |
| 47 | *Peribacillus simplex* | D9_B_73 |
| 48 | *Bacillus* sp. | DX1.1 |
| 49 | *Bacillus hominis* | DX2.1 |
| 50 | *Bacillus hominis* | DX2.3 |
| 51 | *Bacillus* sp. | DX3.1 |
| 52 | *Bacillus* sp. | DX4.1 |
| 53 | *Bacillus velezensis* | DY26 |
| 54 | *Neobacillus* sp*.* | DY30 |
| 55 | *Peribacillus simplex* | E1_1 |
| 56 | *Peribacillus frigoritolerans* | G1S1 |
| 57 | *Bacillus subtilis* | G2S2 |
| 58 | *Paenibacillus* sp. | G2S3 |
| 59 | *Lysinibacillus* sp. | G4S2 |
| 60 | *Bacillus mycoides* | G5S2 |
| 61 | *Bacillus altitudinis* | G6S2 |
| 62 | *Bacillus safensis* | G6S3 |
| 63 | *Bacillus toyonensis* | G7S3 |
| 64 | *Bacillus pumilus* | (GT4_IS1) AKRM3 |
| 65 | *Bacillus altitudinis* | J6-1 |
| 66 | *Bacillus altitudinis* | J6-2 |
| 67 | *Neobacillus cucumis* | JX25 |
| 68 | *Bacillus halotolerans* | KF17 |
| 69 | *Bacillus subtilis* | KF24 |
| 70 | *Rossellomorea marisflavi* | LC18 |
| 71 | *Peribacillus frigoritolerans* | LC22 |
| 72 | *Bacillus wiedmannii* | LN15 |
| 73 | *Bacillus halotolerans* | LN2 |
| 74 | *Peribacillus frigoritolerans* | LN4 |
| 75 | *Cytobacillus firmus* | LN5 |
| 76 | *Lysinibacillus pakistanensis* | LY1 |
| 77 | *Lysinibacillus pakistanensis* | LY18 |
| 78 | *Paenibacillus silvae* | LY60 |
| 79 | *Lysinibacillus pakistanensis* | LY92 |
| 80 | *Bacillus velezensis* | MB7_B11 |
| 81 | *Bacillus velezensis* | MB7_B13 |
| 82 | *Bacillus subtilis* | MB9_B8 |
| 83 | *Bacillus bombysepticus* | (Mi106 D2 head1 chi) AKRM7 |
| 84 | *Bacillus anthracis* | (Mn106-1 head2 chi) AKRM8 |
| 85 | *Bacillus pumilus* | Monterrey_S2 |
| 86 | *Bacillus toyonensis* | Monterrey_S3 |
| 87 | *Bacillus thuringiensis* | Monterrey_S4 |
| 88 | *Bacillus wiedmannii* | Munchenroda_S7 |
| 89 | *Bacillus subtilis* | (MW2 1S1) AKRM4 |
| 90 | *Peribacillus* sp. | NJ11 |
| 91 | *Cytobacillus* sp. | NJ13 |
| 92 | *Bacillus velezensis* | NJ26 |
| 93 | *Peribacillus* sp. | NJ4 |
| 94 | *Bacillus toyonensis* | Puebla_S2 |
| 95 | *Fredinandcohnia* sp. | QZ13 |
| 96 | *Peribacillus simplex* | RZ14 |
| 97 | *Cytobacillus kochii* | RZ2 |
| 98 | *Bacillus pseudomycoides* | SIN1.1 |
| 99 | *Bacillus cereus* | SIN1.2 |
| 100 | *Bacillus hominis* | SIN2.1 |
| 101 | *Bacillus mycoides* | SIN2.2 |
| 102 | *Bacillus mycoides* | SIN2.3 |
| 103 | *Bacillus mycoides* | SIN3.1 |
| 104 | *Bacillus mycoides* | SIN3.2 |
| 105 | *Cytobacillus firmus* | SQ11 |
| 106 | *Neobacillus* sp. | SuZ13 |
| 107 | *Bacillus albus* | SXL388 |
| 108 | *Bacillus halotolerans* | Tehuacan_S4 |
| 109 | *Bacillus cereus* | Tehuacan_S5 |
| 110 | *Fictibacillus enclensis* | TL11 |
| 111 | *Fictibacillus enclensis* | TL8 |
| 112 | *Bacillus velezensis* | TZ19 |
| 113 | *Neobacillus* sp. | WH10 |
| 114 | *Neobacillus cucumis* | WH12 |
| 115 | *Peribacillus simplex* | WH6 |
| 116 | *Siminovitchia fortis* | XLM16 |
| 117 | *Neobacillus novalis* | XLM17 |
| 118 | *Bacillus sonorensis* | YX13 |
| 119 | *Bacillus arachidis* | YX15 |
| 120 | *Neobacillus* sp. | YX16 |
| 121 | *Bacillus cereus* | YX23 |

**Figure S1. Ring-shaped biofilm formation along the inner wall of shake flasks.** The left flask was inoculated with the 121-strain *Bacillales* library, whereas the right flask contained uninoculated medium as a control.

**Figure S2. Biofilm formation (crystal violet staining, A) and lignin-like dye decolorization (B and C) of 15 *Bacillales* strains used as a starting point for SynCom design.** (A) Biofilm formation of 15 *Bacillales* strains monitored with crystal violet staining in a 96 well plate assay. (B) Decolorization of methylene blue (MB; structure shown in inset of panel B graph) of 15 *Bacillales* strains. (C) Decolorization of Azure B (AB; structure shown in inset of panel C graph) of 15 *Bacillales* strains. Data are shown as mean ± SD of three technical replicates.

**Figure S3. Different LDSynCom members exhibit distinct capacities for ring-shaped biofilm formation along the inner wall of shake flasks.** From left to right: *B. paralicheniformis* D9_B_45, *B. paralicheniformis* CEW_1W, and *Paenibacillus* sp. 11B.

**Figure S4. Heatmap representation of top 50 different expressed proteins under M9 and M9+lignin conditions.** Missing were treated as left-censored and imputed using sample-wise minimum log2 intensity minus one prior to z-score transformation.

**Figure S5. Time-course changes in UV absorbance at 280 nm (A280) during shake-flask assay.**
